# Connecting Hippo pathway and cytoophidia in *Drosophila* posterior follicle cells

**DOI:** 10.1101/2023.12.21.572733

**Authors:** Rui-Yu Weng, Lei Zhang, Ji-Long Liu

## Abstract

CTP synthase (CTPS), the rate-limiting enzyme in *de novo* synthesis of CTP, assembles into filamentous structures termed cytoophidia. Here we study the impact of Hippo pathway on the posterior follicle cells (PFCs) in *Drosophila* egg chambers. We find that the inactivation of Hippo pathway correlates with a reduction in cytoophidium length and number within PFCs. During the overexpression of CTPS, the presence of Hippo mutations also reduces the length of cytoophidia in PFCs. In addition, we observe that knocking down CTPS mitigates *hpo* (Hippo)-associated over-proliferation. In summary, our results suggest a connection between the Hippo pathway and the nucleotide biosynthesis enzyme CTPS in PFCs.

## Introduction

Nucleotide metabolism plays a crucial role in various cellular processes, including the synthesis of DNA, RNA and the generation of cellular energy. The synthesis of purines and pyrimidines follows two primary pathways, the *de novo* nucleotide synthesis and the salvage pathway. CTP synthase (CTPS) assumes the role of the rate-limiting enzyme in the *de novo* synthesis pathway for CTP[1, 2]. CTPS exhibits a dynamic equilibrium among monomers, dimers and tetramers[3]. In the present of substrate, CTPS presents at high concentration of active tetramers type[4]. The activity of CTPS is increased in cancer cells, such as liver neoplasia and kidney tumors[5].

In addition to the conventional catalytic activity, CTPS was discovered to form filamentous structures in the 2010, known as cytoophidia[6]. The cytoophidia structures are highly conserved in various species, including *S. cerevisiae*[7], *S. pombe*[8] and mammalian cells[9]. Furthermore, the formation of cytoophidia is regulated by *Myc*[10], *Ras*[11], mTOR-S6K1 pathway[12, 13] and nutritional conditions[14] in the *Drosophila*. Cytoophidia also appear in the cancer, such as human hepatocellular carcinoma[15].

Cytoophida are widely distributed during both the larval stage[16] and oogenesis of *Drosophila*. Within the *Drosophila* ovaries, cytoophidia are observed in various cell types, including follicle cells, nurse cells, and oocytes[17–19]. Cytoophidia can be categorized into two distinct classes: micro-cytoophidia, measuring approximately 1-4 μm in length, and macro-cytoophidia, which can extend up to 30-40 μm[6]. Functionally, cytoophidia serve as scaffolds[20–22] and effectively prolong the half-life of CTPS[23]. Considering that CTPS is the rate-limiting enzyme, it is crucial to gain a comprehensive understanding of its regulatory mechanism.

The Hippo pathway was initially discovered in the *Drosophila* in 1995[24, 25]. The mutations of the Hippo pathway result in over-proliferative phenotype. In *Drosophila*, the core of the Hippo pathway coordinates of a series of phosphorylation events that ultimately inhibit the transcriptional coactivator Yorkie (Yki). Initially, the Sterile 20-like kinase Hippo (Hpo)[26] associates with the WW domain scaffolding protein Salvador (Sav)[27] to phosphorylate and activate the DBF family kinase Warts (Wts). Subsequently, the activated Wts interacts with Mob as tumor suppressor (Mats)[28] to phosphorylate and inhibit Yki[29], creating a 14-3-3 binding site. Yki then binds with 14-3-3 protein, retaining Yki in the cytoplasm[30].

The Hippo pathway is evolutionarily conserved in the mammals[31]. The Hippo pathway controls the expression of downstream genes, including *diap1*, *bantam* and *cyclin E*[32] to regulate proliferation and apoptosis, thereby maintaining tissue size homeostasis[33]. Furthermore, the Hippo pathway is implicated in the regulation of lipid metabolism and organ regeneration [34]. Dysregulation of the Hippo pathway has been identified in various cancers, such as malignant mesothelioma[35] and prostate adenocarcinoma[36]. Therefore, our study aims to investigate the connection between the Hippo pathway and cytoophidia.

Using the *Drosophila* egg chamber as a model[37], we observe that Hippo pathway and cytoophidium assembly interconnect in posterior follicle cells (PFCs). Mutations in Hippo pathway result in over-proliferation and Notch accumulation in the PFCs. Hippo pathway mutations reduce the length and number of cytoophidia within the PFCs. In addition, Hippo pathway mutations reduces the length of cytoophidium, even when CTPS is overexpressed. CTPS knockdown suppresses the over-proliferative phenotype in *hpo*-mutant PFCs. Therefore, our study connects the Hippo pathway and cytoophidia in PFCs.

## Results

### Hippo pathway is essential for oocyte polarity

In *Drosophila* oogenesis, the establishment of the body axis depends on the polarization of the oocyte [38, 39]. The asymmetric distribution mRNA and proteins within the oocyte plays a pivotal role in determining the anterior-posterior (AP) body axis[40]. For example, *gurken* (*grk*) mRNA localizes adjacent to the oocyte nucleus[41], while *bicoid* (*bcd*) mRNA is found at the anterior pole[39], and *osker* (*osk*) mRNA is concentrated at the posterior pole[38]. Dysregulation of Hippo pathway disrupts oocyte migration, leading to over-proliferation of anterior and posterior follicle cells, as well as mislocalization of Osk and Staufen (Stau)[42, 43]. In this study, we aim to investigate the phenotypic consequences in posterior follicle cells when the Hippo pathway undergoes mutations.

We used mosaic analysis with a repressible cell marker (MARCM) technique to generate FRT/FLP mitotic clones of mutated cells with a loss of function (*wts^x1^*) allele of *wts*. In oocytes, the OSK signal localizes in the crescent at the posterior side (Fig 1A and B), while single-layer follicle cells surround germline cells (Fig 1C). The Notch intracellular domain (NICD) signal was observed at the apical part of the follicle cells (Fig 1A and D). Our findings reveal that the OSK mis-localizes to the middle of the oocyte (Fig 1F and G) when *wts* mutates in PFCs. An over-proliferative phenotype was observed in the *wts^x1^* PFCs (Fig 1F and H). When the posterior follicle cells are GFP-positive clones, Notch signals accumulate (Fig 1I and J). These results are consistent with previous studies [42, 43].

**Fig 1.**
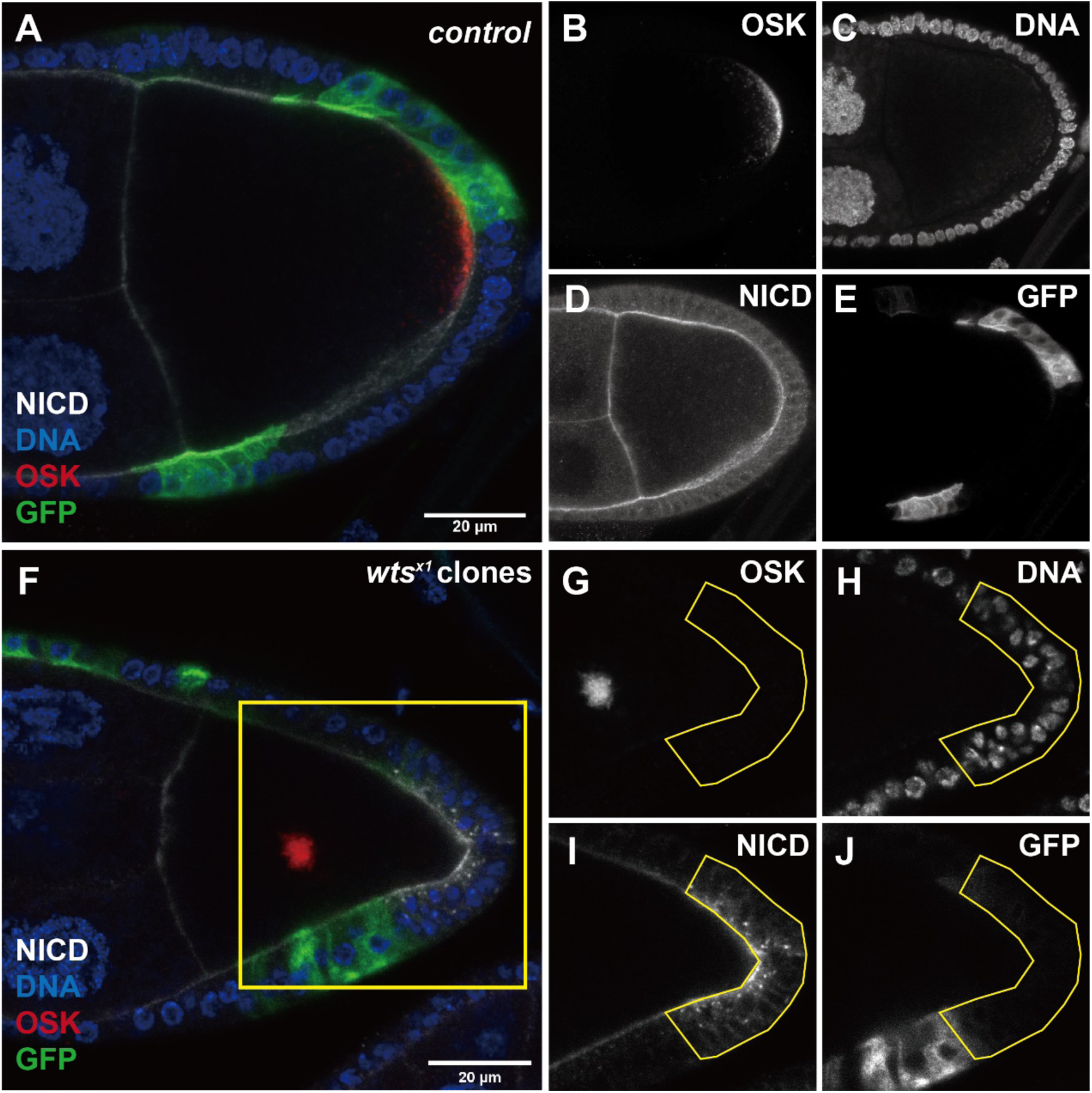
Wts is required for oocyte polarity. (A-E) The Oskar (OSK, red in A, white in B) protein localizes at the posterior crescent of the stage-9 FRT82B control oocyte. Notch intercellular domain (NICD, white in A and D) labels the Notch signal, which is located in the membrane of the follicle cells and germline cells. (F-J) When *wts* mutates in posterior follicle cells, OSK (red in F, white in G) is relocated to the middle of the stage-9 oocyte. The *wts* mutant posterior follicle cells show an over-proliferative phenotype and a disordered distribution of NICD (white in F and I). DNA is stained with Hoechst 33342 (blue in F, white in H). Recombinant cells are marked with GFP (green in F, white in J). Scale bars, 20 μm.

### Cytoophidium length and number are decreased in *wts*-mutant PFCs

Hippo pathway regulates the expression of downstream genes (*bantam*, *cyclin E* and *diap1*) to control cell proliferation and apoptosis. Previous studies have revealed the connections between the Hippo pathway and various metabolic pathways, such as glucose metabolism, amino acid metabolism, and lipid metabolism[34]. Additionally, the Hippo pathway receives upstream signals from components such as the apical complex crumbs (Crb) and basolateral complex Scrib/Dlg/Lgl[44–46].

Mutations in the Hippo pathway can affect cell polarity and cell adhesion, such as *hpo*, *wts* and *yki*[43]. Importantly, cytoophidia are predominantly localized at the basolateral side of follicle cells[21]. Furthermore, it is known that Yki can regulate the expression of *Myc*[47]. Our previous study found the essential role of *Myc* in cytoophidia assembly[10]. Therefore, our current investigation aims to elucidate the potential involvement of the Hippo pathway in cytoophidium assembly.

We observed that the *wts^x1^* mutation induced over-proliferative phenotype (Fig 1H) and resulted in the accumulation of NICD signals in PFCs (Fig 1I). We further if the *wts^x1^* mutation affects cytoophidia assembly in PFCs. We used antibodies targeting the NICD and CTPS to perform immunostaining of egg chambers. Cytoophidia were observed in germline cells and follicle cells (Fig 2A and D). The NICD signal localizes at apical side of follicle cells (Fig 2B) and one layer of follicle cells surround the germline cells (Fig 2C). The over-proliferative phenotype appears in *wts^x1^*PFCs (Fig 2F and H). Based on the accumulation of NICD signals (Fig 2G), we quantified the length of cytoophidia in follicle cells (Fig 2I). Analysis revealed that the length of cytoophidia was reduced compared to adjacent cells (Fig 2K). Counting the number of cytoophidia and follicle cells, we found that the proportion of cytoophidium in NICD accumulated posterior follicle cells is decreased. There are no different between other GFP-positive clones and GFP-negative clones (Fig 2L).

**Fig 2.**
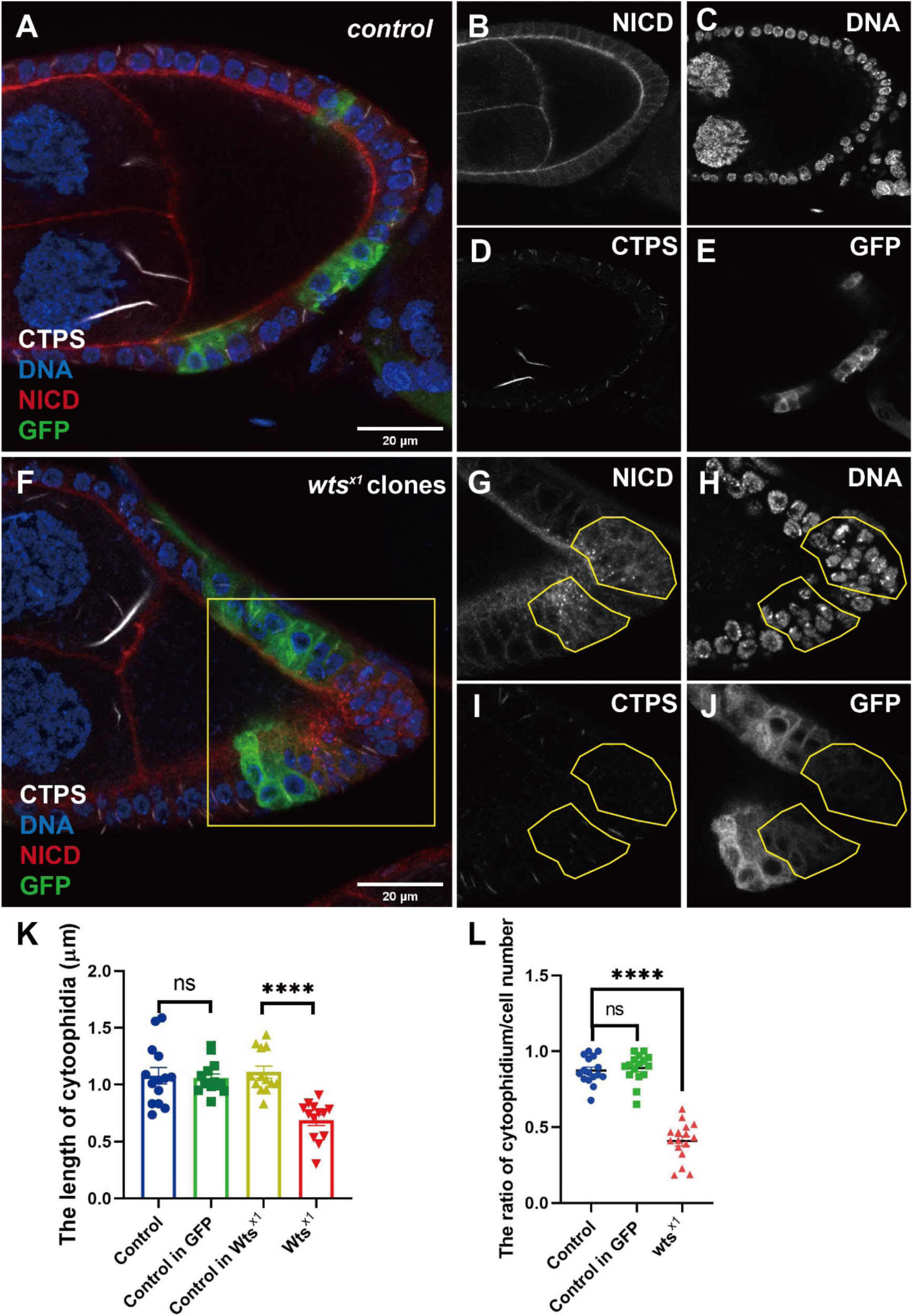
*Wts^x1^* decreases cytoophidium length and number in posterior follicle cells. (A-E) In the FRT82B control egg chamber, there is no difference in the NICD signal (red in A, white in B) and the nuclear size (blue in A, white in C) of follicle cells between GFP-positive (green in A, white in E) and GFP-negative clones. Cytoophidia labeled with CTPS (white in A and D) exist in follicle cells and germline cells. There is no difference in cytoophidium length between GFP-positive and GFP-negative clones. (F-J) In the *wts^x1^* egg chamber, NICD (red in F, white in G) accumulates in posterior follicle cells (yellow box in F). After *wts* mutation, the size of the cell nucleus and the length of cytoophidia in posterior follicle cells decrease, while the number of cells increases. DNA is stained with Hoechst 33342 (blue in F, white in H). Recombinant cells are marked with GFP (green in F, white in J). Scale bars, 20 μm. (K) Quantification shows that *wts^x1^* decreases the cytoophidium length in posterior follicle cells. ****P<0.0001, n.s. = no significant, mean with SED (Student’s *t* test). (L) Quantification shows that *wts^x1^*decreases the number of cytoophidium in posterior follicle cells. ****P<0.0001, n.s. = no significant, mean with SED (Student’s *t* test).

### Cytoophidium length and number are decreased in *hpo*-mutant PFCs

To confirm that the reduction in cytoophidium length can indeed be regulates by the Hippo pathway, we generated FLP/FRT mitotic clones in the egg chamber using a truncating (*hpo^BF33^*) allele of *hpo*. The NICD signal and nucleus size have no significant differences between GFP-positive and GFP-negative clones in control egg chambers (Fig 3A-C). The length of cytoophidium has no significant differences between GFP-positive and GFP-negative clones (Fig 3D and E). The signal of NICD also accumulates in the *hpo* mutant posterior follicle cells (Fig 3F and G).

**Fig 3.**
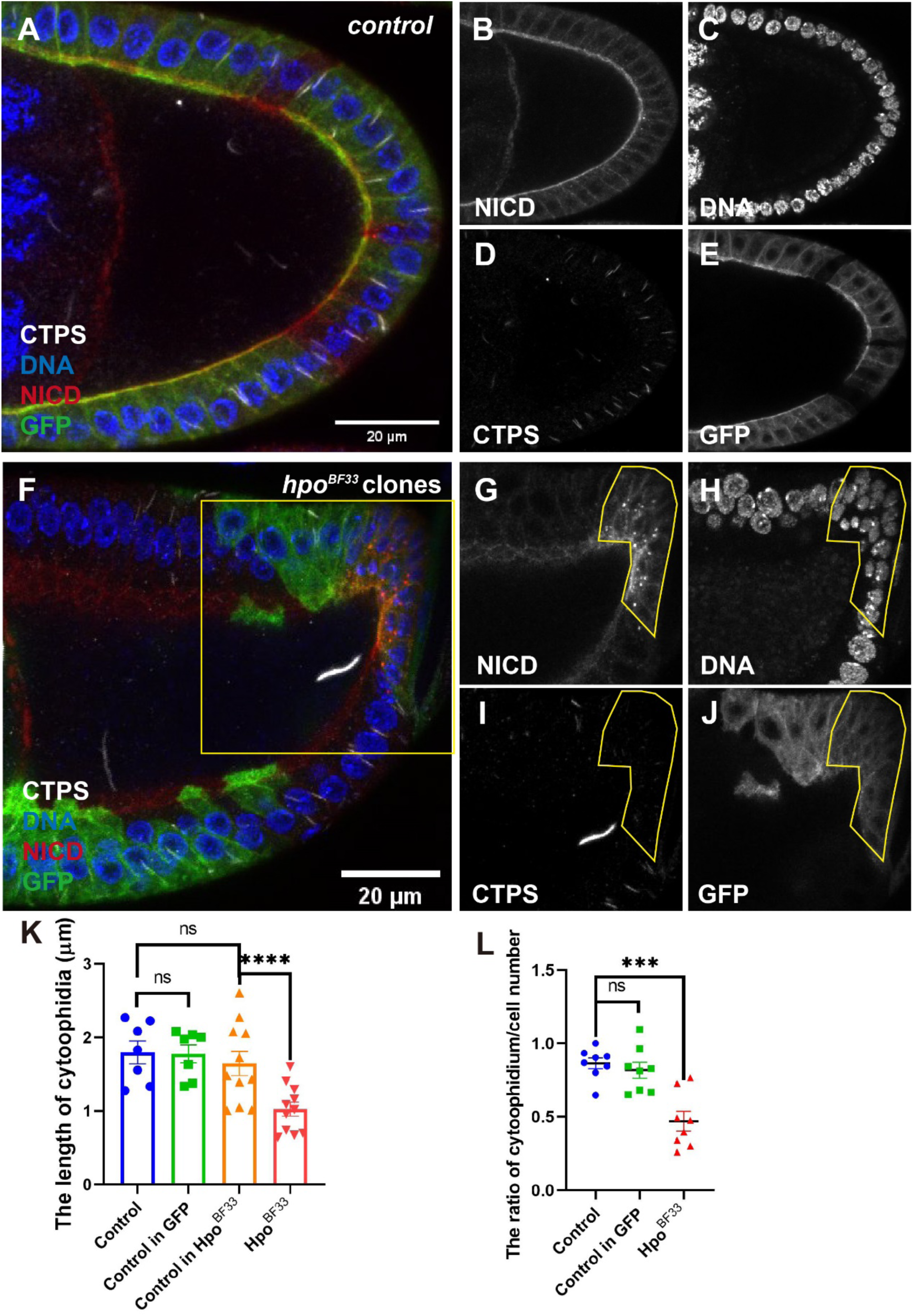
*Hpo^BF33^* decreases cytoophidium length and number in posterior follicle cells. (A-E) In the FRT42D control egg chamber, there is no difference in the NICD signal (red in A, white in B) and the nuclear size (blue in A, white in C) of follicle cells between GFP-positive (green in A, white in E) and GFP-negative clones. Cytoophidia labeled with CTPS (white in A and D) exist in follicle cells and germline cells. There is no difference in cytoophidium length between GFP-positive and GFP-negative clones. (F-J) In the *hpo^BF33^* egg chamber, NICD (red in F, white in G) accumulates in posterior follicle cells (yellow box in F). After *hpo* mutation, the size of the cell nucleus and the length of cytoophidia in posterior follicle cells decrease, while the number of cells increases. DNA is stained with Hoechst 33342 (blue is F, white in H). Recombinant cells are marked with GFP (green in F, white in J). Scale bars, 20 μm. (K) Quantification shows that the *hpo* mutant decreases the cytoophidium length in posterior follicle cells. ****P<0.0001, n.s. = no significant, mean with SED (Student’s *t* test). (L) Quantification shows that *hpo^BF33^*decreases the number of cytoophidium in posterior follicle cells. ***P<0.001, n.s. = no significant, mean with SED (Student’s *t* test).

The over-proliferative phenotype is also observed in the NICD-accumulated posterior follicle cells which are marked by yellow frame (Fig 3H). Furthermore, when we quantified the length of cytoophidia, we observed that the *hpo* mutation also resulted in a decrease in cytoophidium length in PFCs (Fig 3I and K). Similarly, the proportion of cytoophidium in *hpo* mutant posterior follicle cells is decreased. There are no different between other GFP-positive clones and GFP-negative clones (Fig 3L)

### *Hpo* mutation decreases cytoophidium length even if CTPS is overexpressed

The length of the cytoophidium is directly related to the concentration of intracellular CTPS. It is known that overexpression of CTPS increases the size and abundance of cytoophidia[14, 18, 48]. Acting as a scaffold, CTPS overexpression can also lead to the thickening of follicle cells[21]. To investigate whether CTPS overexpression can rescue cytoophidia length in the *hpo* mutant PFCs, we generated a stock containing the *hpo^BF33^* mutation and *UAS-CTPS-mCherry*. The signal of NICD was used as a marker. The results revealed an increase in the length and abundance of cytoophidia within the GFP-positive clones (Fig 4A and D).

**Fig 4.**
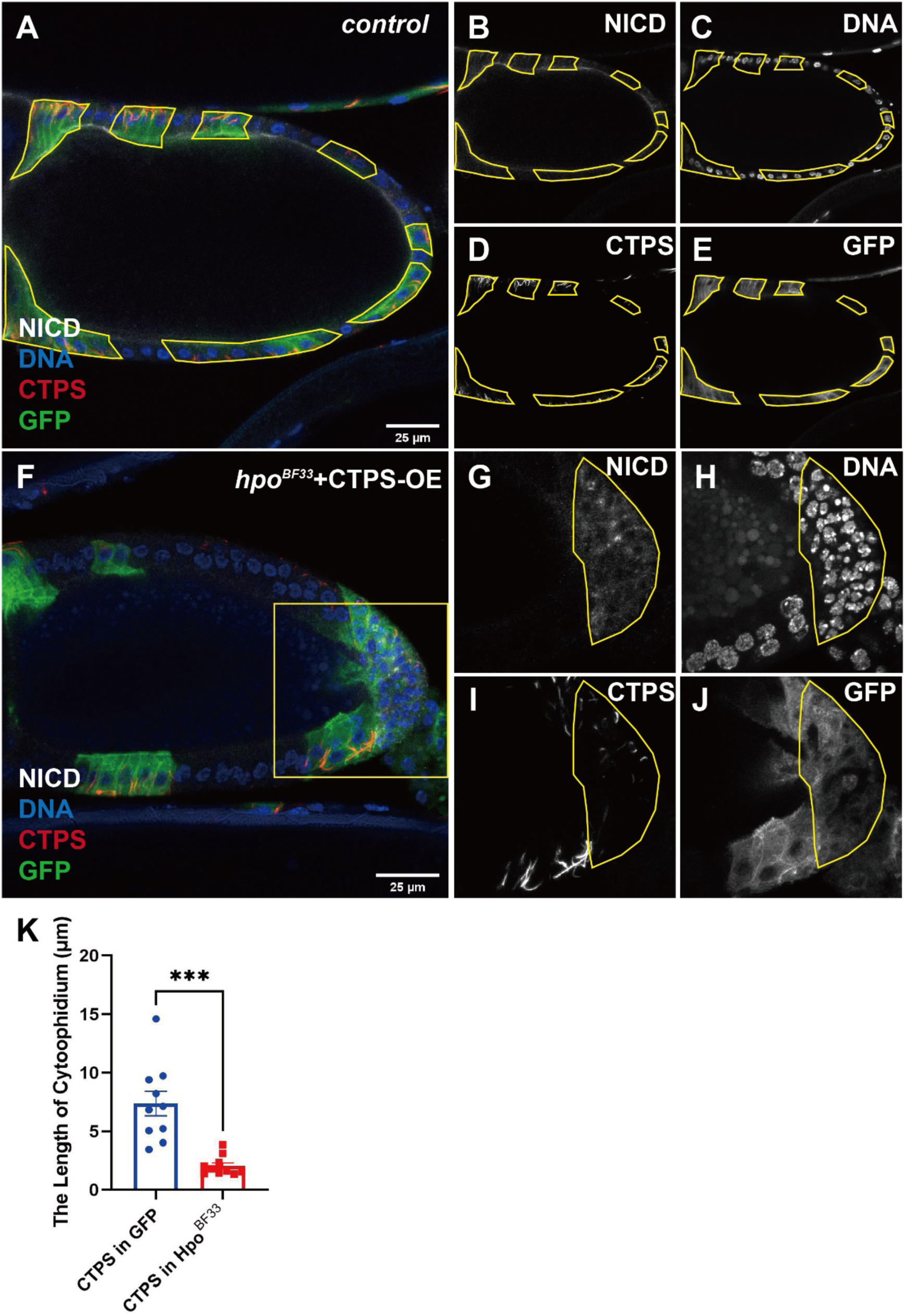
Even if CTPS is overexpressed, the *hpo^BF33^* decreases cytoophidium length. (A-E) In the FRT42D with CTPS overexpression egg chamber, there is no difference in the NICD signal (white in A and B) of follicle cells between GFP-positive (green in A, white in E) and GFP-negative clones. The length and number of cytoophidia labeled with CTPS (red in A, white in D) increase in the GFP-positive follicle cells. DNA is stained with Hoechst 33342 (blue is A, white in C). (F-J) In the *hpo* mutant with CTPS overexpression, NICD (white in F and G) accumulates in posterior follicle cells (yellow box in F). After *hpo* mutation, the length of cytoophidium with overexpressed CTPS in posterior follicle cells (yellow box in I) is shorter than that in other GFP-positive regions. DNA is stained with Hoechst 33342 (blue in F, white in H). Recombinant cells are marked with GFP (green in F, white in J). Scale bars, 25 μm. (K) Quantification shows *hpo* mutation decreased the cytoophidium length in posterior follicle cells, even if CTPS is overexpressed. ***P<0.001, mean with SED (Student’s *t* test).

No significant differences were observed in NICD signal and cell nucleus size between GFP-positive and GFP-negative clones in control egg chambers (Fig 4B and C). Notably, we observed an increase in cytoophidium length within the GFP-positive clones (Fig 4F and I) in *hpo^BF33^* mutant follicle cells. Additionally, NICD signal accumulation and over-proliferative phenotype were observed in the *hpo*-mutant PFCs (Fig 4G and H) which are marked by yellow frame. It is worth noting that we found that the length of cytoophidia in NICD signal accumulation clones was shorter than that in other GFP-positive clones (Fig 4I and K).

### CTPS knockdown suppresses *hpo^BF33^* induced excessive proliferation

Previous found that CTPS knockdown could suppress *Myc*-induced overgrowth phenotype[11]. Hence, we aim to investigate whether the CTPS is required for the Hippo pathway activity. To answer this question, we generated a stock containing *hpo^BF33^* and *UAS-CTPS-RNAi*. The over-proliferative phenotype and NICD-accumulated phenotype appear in the *hpo^BF33^* PFCs (Fig 5A-A’’’). Our observations demonstrate the absence of cytoophidium in the GFP-positive clones (Fig 5B). Surprisingly, we also observe the NICD signal is accumulated in the GFP-positive PFCs (Fig 5B-B’’’). To analyze the volume of NICD accumulated signal, we constructed a surface using Imaris within the NICD accumulated signal. Quantification of the volume of NICD accumulated signal suggests that phenotype in the *hpo^BF33^* with CTPS knockdown is smaller than *hpo^BF33^*only egg chamber (Fig 5E). These results support the potential role of CTPS within the Hippo pathway.

**Fig 5.**
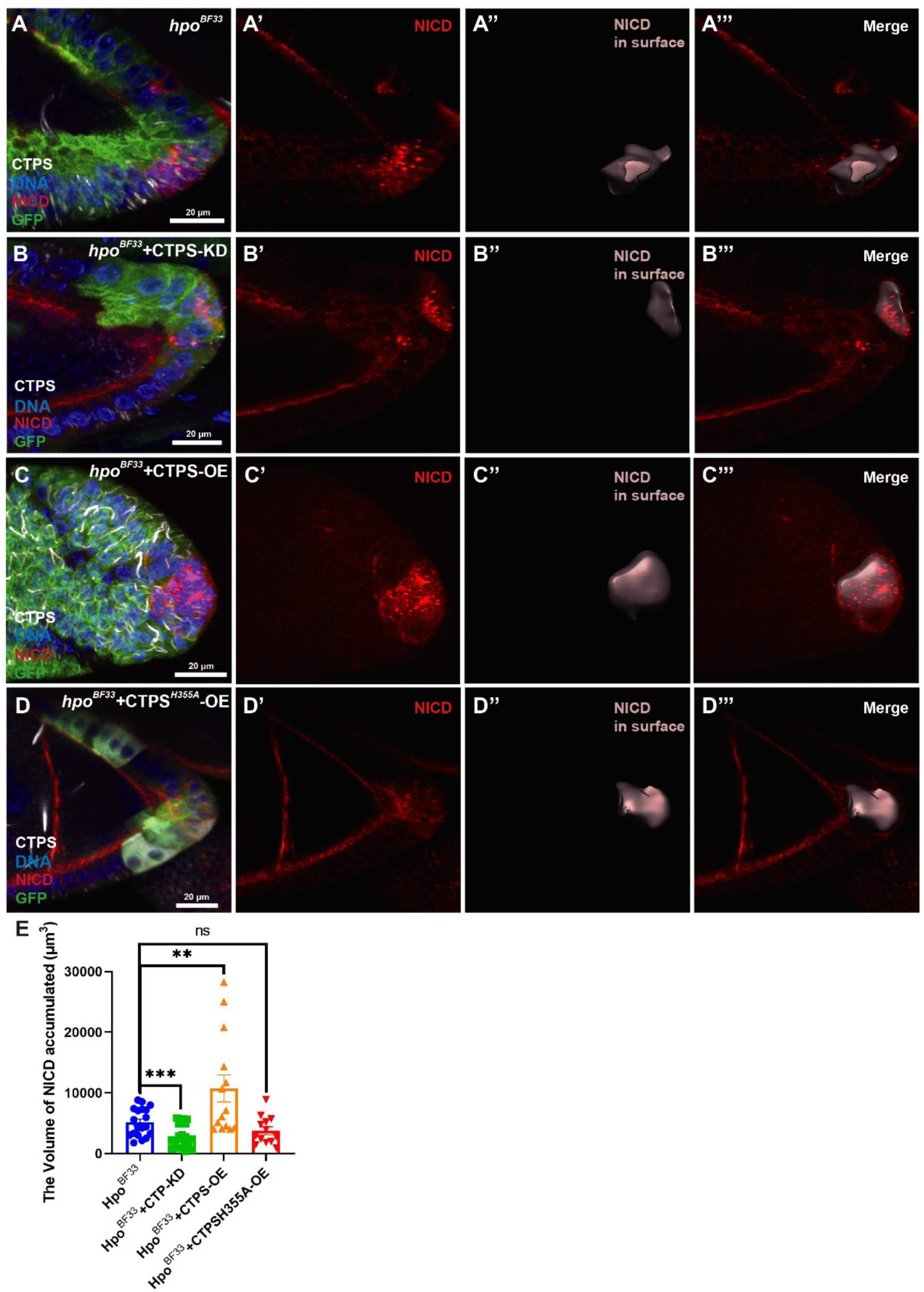
CTPS plays a role in the over-proliferative phenotype in *hpo* mutant PFCs. (A-A’’’) In the 3D view of *hpo* mutant egg chamber, NICD (red in A and A’) accumulates in posterior follicle cells (light red box in A’’ and A’’’). The merge image shows the surface within the NICD accumulated signal. (B-B’’’) In the 3D view of *hpo* mutant with CTPS knockdown, NICD (red in B and B’) accumulates in posterior follicle cells (light red in B’’ and B’’’). The merge image shows the surface within the NICD accumulated signal. (C-C’’’) In the 3D view of *hpo* mutant with CTPS overexpression, NICD (red in C and C’) accumulates in posterior follicle cells (light red in C’’ and C’’’). The merge image shows the surface within the NICD accumulated signal. (D-D’’’) In the 3D view of *hpo* mutant with *CTPS^H355A^* overexpression, NICD (red in D and D’) accumulates in posterior follicle cells (light red in D’’ and D’’’). The merge image shows the surface within the NICD accumulated signal. Scale bars, 20 μm. (I) Quantification shows that CTPS overexpression promotes the volume of over-proliferative phenotype. CTPS knockdown inhibits the volume of over-proliferative phenotype in *hpo*-mutant PFCs. CTPS^H355A^ overexpression has no difference with *hpo-*mutant egg chamber. **P<0.01, ***P<0.001, n.s. = no significant, mean with SED (Student’s *t* test).

### CTPS overexpression increases the volume of NICD in *hpo* mutant PFCs

Increased CTPS activity is a characteristic of many forms of cancer, such as hepatomas and leukemia[5, 49, 50]. Although the *hpo^BF33^* mutation still resulted in a decrease in cytoophidium length in PFCs, we attempted to determine whether CTPS overexpression exacerbates over-proliferation associated with *hpo^BF33^*. In the *hpo^BF33^*with CTPS overexpression egg chamber, the volume of NICD accumulation is marked in a light red box (Fig 5C-C’’’). Quantification showed that CTPS overexpression increased the volume of NICD accumulated in *hpo^BF33^*PFCs (Fig 5E).

### *CTPS^H355A^* overexpression does not increase the volume of NICD in *hpo* mutant PFCs

In order to investigate if the cytoophidium structure affect the proliferation controlled by Hippo pathway, we generated a stock containing *hpo^BF33^* and *UAS-CTPS^H355A^-mCherry-HA*. It is worth noting that the cytoophidium is depolymerized when the His355 is replaced by Alanine. The H355A mutation only affects polymerization, but not the enzymatic activity[22, 51, 52]. While the CTPS^H355A^ overexpression was observed in the GFP-positive clones (Fig 5D), the NICD signal was found to accumulate in the *hpo*-mutant PFCs (Fig 5D-D’’’). Notably, CTPS^H355A^ overexpression showed no significant difference with *hpo^BF33^* only PFCs in the volume of NICD-accumulated signal (Fig 5E). These results suggest that cytoophidium structure contributes to the over-proliferative phenotype induced by the Hippo pathway.

## Discussion

Cancer and metabolism have long been at the forefront of research. In tumor cells, cell metabolism is accelerated, leading to excessive cell proliferation[53]. The Hippo pathway plays a crucial regulatory role in cell metabolism, such as glucose metabolism, glutamine metabolism and lipid metabolism[34, 54]. The dysregulation of Hippo pathway is observed in various human cancers, such as breast cancer[55], glioblastoma[56], hepatocellular carcinoma and cholangiocarcinoma[57]. Overexpression of YAP results in massive liver size compared to normal size[58, 59].

The interaction of YAP/TAZ with TEAD is considered a potential druggable site. However, it has been reported that the small molecule verteporfin, which inhibits YAP-induced liver growth, also induces cell death in normal cell[60]. Previous study found that YAP1 increases glutamine level and enhances *de novo* nucleotide synthesis in zebrafish[54, 61]. Knockdown of YAP/TAZ decreased the expression of CTP synthase 1 in MDA-MB-231 cell[62]. However, the mechanism by which the Hippo pathway regulates nucleotide synthesis remains unknown.

The CTPS activity and levels are increased in cancer cells such as hepatomas[5, 50]. The length of cytoophidia is related with the concentration of CTPS in the cell[63]. In mice, inhibiting CTPS1 suppresses tumor growth and induces apoptosis[64], indicating that CTPS plays a role in tumor metabolism.

The formation of cytoophidia regulates enzyme activity. For example, cytoophidia formed by inosine monophosphate dehydrogenase (IMPDH) reflect the upregulation of purine synthesis in mammalian cell lines[65]. Filamentation of human CTPS increases enzymatic activity in vitro[66]. However, CTPS cytoophidia inhibit enzymatic activity in *S. cerevisiae*, *E. coli* and *D. melanogaster*[9, 48, 67, 68]. CTPS is considered a potential drug target[64]. The glutamine analog 6-Diazo-5-oxo-L-norleucine (DON) binds to the glutamine binding site in the CTPS GAT domain, inhibiting CTPS activity and increasing cytoophidia length[17, 69].

The Hippo pathway regulates cell proliferation and apoptosis. Overexpression of Yki increases the intestinal stem cell proliferation[70, 71]. In the *Drosophila* follicle cells, mutations of Hippo pathway induce over-proliferation in the posterior follicle cells. We found that mutations of *wts* and *hpo* decrease the length and number of cytoophidium in PFCs. During the stage 10a-b of oogenesis, the follicle cells undergo rapidly growth and cytoophidia are decreased[10]. Hence, we speculate that the CTPS cytoophidia may depolymerize to release more active CTPS for CTP synthesis in hippo-mutant PFCs.

Overexpression of CTPS promotes cytoophidium assembly and increases CTP concentration[48]. Our study shows that the length of cytoophidium in *hpo*-mutant PFCs is shorter than cytoophidia in neighboring GFP-positive follicle cells and the over-proliferation is promoted when CTPS is overexpressed. Knockdown of CTPS suppresses the over-proliferation induced by Hippo pathway mutation, indicating that CTPS plays a role in the regulation of cell proliferation by Hippo pathway. Modulating the activity of metabolic enzymes may offer potential treatment options for cancer caused by Hippo mutations. Hence, it is possible to use DON to target CTPS to suppress the over-proliferative phenotype induced by Hippo mutations.

In conclusion, our data demonstrate that the Hippo pathway regulates CTPS cytoophidium formation in *Drosophila* posterior follicle cells. Interestingly, when CTPS is overexpressed, the Hippo pathway still decreases the length of cytoophidia. Additionally, knockdown of CTPS suppresses the over-proliferation caused by *hpo^BF33^*. Collectively, this study provides a novel insight into research on the Hippo-related cancer by establishing a connection between the Hippo pathway and CTPS cytoophidia.

## Materials and methods

### Fly Stocks

All stocks were maintained at 25 ℃ on standard cornmeal.

### The MARCM system

*hsFlp; FRT42 tubulin-GAL80, tubulin-GAL4*/Sp; *UAS-GFP*/Tm6B; *yw*,

*hsFlp, UAS-GFP*;

*FRT82B tubulin-GAL80, tubulin-GAL4*/Tm6B;

*yw; FRT42D hpo^BF33^*/Cyo;

Control: *yw; FRT42D*/Cyo;

*yw; FRT82B wts^latsX1^*/Tm6B;

Control: *yw; FRT82B*/Tm6B.

The transgenic RNAi line *UAS-CTPS-RNAi* was obtained from the TsingHua Fly Center, stock number THU2302; The homozygote *UAS-CTPS-mCherry* in the Chromosome III; the homozygote *UAS-CTPS^H355A^-mCherry-HA* in the Chromosome III; If/Cyo; Sb/Tm6B is a gift from *Yuu Kimata’s* lab.

We generate *Hpo^BF33^*/Cyo; *UAS-CTPS-RNAi*/Tm6B by crossing *Hpo^BF33^*/Cyo with double balancer flies for two generations to construct *Hpo^BF33^*/Cyo; Sb/Tm6B. Crossing *UAS-CTPS-RNAi* flies with double balancer flies for two generations to construct If/Cyo; *UAS-CTPS-RNAi*/Tm6B. Then, crossing *Hpo^BF33^*/Cyo; Sb/Tm6B with If/Cyo; *UAS-CTPS-RNAi*/Tm6B to get *Hpo^BF33^*/Cyo; *UAS-CTPS-RNAi*/Tm6B strain. The control strain *FRT42D*/Cyo; *UAS-CTPS-RNAi*/Tm6B is constructed by same method.

The *Hpo^BF33^*/Cyo; *UAS-CTPS-mCherry/Tm6B* and *FRT42D*/Cyo; *UAS-CTPS-mCherry*/Tm6B, *Hpo^BF33^*/Cyo; *UAS-CTPS^H355A^-mCherry-HA/Tm6B* and *FRT42D*/Cyo; *UAS-CTPS^H355A^-mCherry-HA*/Tm6B are constructed by same method.

### Immunohistochemistry

The tissues of female *Drosophila* for 3-5 days were dissected from at least three technical replicates, and each replicate contained more than 5 flies. Mosaic tissues were obtained with the FLP/FRT system with hsFLP drivers. For the loss of function, adult flies were heated shock for 1 hour and 37 °C and heat shock for 1 hour at the third day, dissected at 3 days after heat shock.

Ovaries from flies were dissected in Grace’s Insect Medium (Gibco) and then fixed in 4 % formaldehyde (Sigma), 10 μl 40 % formaldehyde diluted in 90 μl PBS for 10 min before immunofluorescence staining.

The samples were then washed by PST (0.5 % horse serum + 0.3 % Triton X-100 in PBS) and incubation in PST for about 1 hour to block. The samples were incubated in primary antibodies overnight at room temperature. Then, they were briefly rinsed with PST and incubated in secondary antibodies and Hoechst 33342 (1:5,000; Thermo Fisher, Rockford, IL, USA) for DNA staining two hours at room temperature.

Primary antibodies used in this study were rabbit anti-CTPS (1:1000; y-88, sc-134457, Santa Cruz BioTech Ltd., Santa Cruz, CA, USA), mouse anti-Notch (1:100, C17.9C6, Developmental Studies Hybridoma Bank), rabbit anti-Oskar (1:500). Secondary antibodies used in this study were anti-mouse Alexa Fluor® 594 nm (invitrogen A11032), donkey anti-rabbit Cy3 (Jackson 711-165-152), donkey anti-mouse Cy5 (jackson 715-005-151), donkey anti-rabbit Cy5 (jackson 711-175-152).

### Microscopy and Image Analysis

All images were obtained under laser-scanning confocal microscopy (Zeiss 980). Image processing was performed using Zeiss Zen (version 3.4). ImageJ was used to analyze the length of cytoophidium and the number of cytoophidium and cell number (Downland from <u>ImageJ</u>). For each statistical quantification, we collected the images using Zeiss 980 with the interval as 0.5 μm for z-stack by using 40 Х objective, more than 3 egg chambers were quantified per genotype, biological repeats = 3. Representative images with maximum intensity projection are shown in all figures.

## Data Analysis

Images collected by confocal microscopy were processed using Adobe Illustrator, ImageJ and Imaris x64 9.0.1. The surface is built on the basis of segment only a region of interest. Quantitative analysis was processed by Excel and GraphPad Prism 9. For all experiments, the data are represented as average ± SEM. Student’s t-test was performed to check for significant differences between data groups. In dot plots, a dot represents an egg chamber.

## Credit author statement

**Rui-Yu Weng**: Conceptualization, Methodology, Investigation, Formal analysis, Writing – Original Draft, Visualization. **Ji-Long Liu**: Conceptualization, Resources, Writing – Review & Editing, Supervision, Project administration. **Lei Zhang**: Conceptualization, Resources, Writing – Review & Editing, Supervision, Project administration.

## Acknowledgments

We thank the Molecular Imaging Core Facility (MICF) at school of Life Science and Technology, ShanghaiTech University for providing technical support. We thank the Core Facility of Drosophila Resource and Technology.

